# A Conserved Locus Coeruleus fMRI Signature of Brain-State Transitions across Sleep, Anesthesia, and Wakefulness

**DOI:** 10.64898/2026.01.28.702261

**Authors:** Francesca Barcellini, Georgios Foustoukos, Daniel Wenz, Brandon R Munn, James M Shine, Laura M.J. Fernandez, Anita Lüthi, Valerio Zerbi

**Author notes:** These authors contributed equally to this work (Francesca Barcellini, Georgios Foustoukos). Shared last authorship (Anita Lüthi, Valerio Zerbi). **Correspondence**: Valerio Zerbi, University of Geneva, Campus Biotech, Chemin des Mines 9 B1.01 – 255.040, 1202, Genève, Switzerland.

## Abstract

Neuromodulatory systems dynamically reconfigure large-scale brain networks to support adaptation across behavioral and cognitive states. The locus coeruleus (LC), which broadcasts noradrenaline throughout the forebrain, is a central regulator of arousal and state-dependent dynamics. However, how LC activity manifests in brain-wide organization across physiological contexts, and how it biases fMRI connectivity, remains poorly understood.

Using an optogenetically informed cross-species framework, we identify a transient LC-derived spatiotemporal pattern of brain activity accompanying brain-state transitions under progressively naturalistic conditions: controlled LC stimulation and endogenous LC fluctuations in anesthetized mice, sleep–wake transitions in rodents and humans, and resting-state activity in awake humans. This LC-derived signature is conserved across species and contexts, leaving a robust and detectable imprint on the BOLD signal. Critically, the prevalence of LC events systematically biases functional connectivity metrics in human fMRI.

These findings establish LC activity as a mechanistically interpretable source of variability in resting-state measurements, with direct implications for the interpretation of fMRI biomarkers in arousal-related disorders.

## Introduction

The brain adapts to ever-changing environments by flexibly reorganizing information flow through its networks. This flexibility depends on the brain’s ability to periodically shift between different functional modes: introspective states, in which brain regions operate more independently, and engaged states, in which regions communicate more broadly across networks to support focused action and learning. This framework of segregation and integration is supported by computational network models, which demonstrate that shifting between these configurations optimizes information processing for different behavioral demands [1], [2], [3]. Yet how these state transitions occur at the circuit level remains unclear. A growing body of evidence implicates neuromodulatory systems as key drivers: by modulating neural gain and reshaping circuit interactions, neuromodulators can dynamically shift the brain between segregated and integrated states [4], [5], [6]. These dynamics have important implications for interpreting both resting-state fMRI (rs-fMRI) signals and the functional connectome. Neuromodulatory influences can bias connectivity patterns toward specific network configurations that are state-dependent, shifting with internal neuromodulatory fluctuations. This distinction is increasingly critical as the field moves toward precision medicine, in which rs-fMRI is used to develop individualized biomarkers, objective measures that predict an individual’s disease status, treatment response, or cognitive profile [6], [7], [8]. Understanding how internal state and neuromodulatory tone shape fMRI signals is thus essential for distinguishing trait-like neural signatures from state-dependent fluctuations in functional connectivity.

Among the brain’s neuromodulatory systems, the locus coeruleus (LC), a small noradrenergic brainstem nucleus, stands out as a key regulator of brain states. The LC projects extensively throughout the forebrain and operates across diverse behavioral contexts: it responds to salient stimuli, maintains attention during wakefulness, and facilitates sleep-stage transitions [10], [11], [12]. These functions are expressed through distinct firing patterns and are mediated by different postsynaptic adrenergic receptor signaling mechanisms, which shape neural gain and circuit dynamics across cortical and subcortical targets [13], [14]. During wakefulness, tonic LC activity sustains arousal and promotes network integration, whereas phasic bursts modulate attention to salient events. During sleep, LC activity undergoes structured changes that reflect transitions between sleep stages [15], [16], [17]. Recent preclinical optogenetic and chemogenetic studies have demonstrated that distinct LC firing patterns reliably produce large-scale changes in brain network organization, fundamentally reshaping fMRI signals across widespread cortical and subcortical regions [18], [19], [20]. Determining whether these LC-induced network changes also arise during natural state transitions in humans is therefore an important open question, as their presence would imply that LC activity leaves measurable traces in the resting-state connectivity observed in clinical fMRI. Yet it remains unknown whether such LC-driven signature emerges in humans and how it shapes the fMRI signal.

In this work, we describe a stereotypical fMRI signature that LC activity imprints on blood-oxygen-level-dependent (BOLD) signal, and its impact on rs-fMRI functional connectivity. We introduce an optogenetically-informed, cross-species framework to ask whether LC-evoked specific BOLD patterns recur under increasingly naturalistic conditions, ranging from controlled stimulation, endogenous LC fluctuations in anesthetized mice, sleep-wake transitions in rodents, analogous transitions in humans, and phasic events during awake human rest. We demonstrate that the LC-derived signature is indeed conserved across all these contexts, revealing how LC activity fundamentally shapes resting-state connectivity. Moreover, we show that the strength of functional connectivity measured in human rs-fMRI is directly influenced by the overall strength (root mean square, RMS) of LC-signature expression during the scan. These findings establish that endogenous LC fluctuations leave measurable imprints on functional connectivity and represent a potential biomarker of arousal and state regulation, with direct implications for interpreting resting-state patterns in psychiatric and neurological conditions.

## Results

### 1. Optogenetic stimulation maps a stereotyped LC fMRI signature

To establish a reference LC fMRI signature under controlled conditions, we reanalyzed the optogenetic–fMRI dataset from Grimm et al. [18], in which mice expressing channelrhodopsin-2 in LC neurons underwent blue-light stimulation at 3, 5, or 15 Hz under light isoflurane anesthesia (1.1% isoflurane; 9 cycles of 30-s ON/OFF per session; N = 15 - 18 per frequency; N = 32 sham controls; Fig 1A). While the original study emphasized frequency-dependent effects, we focused instead on responses that remained invariant across all stimulation frequencies, aiming to extract the fundamental, context-independent LC network architecture.

**Figure 1.**
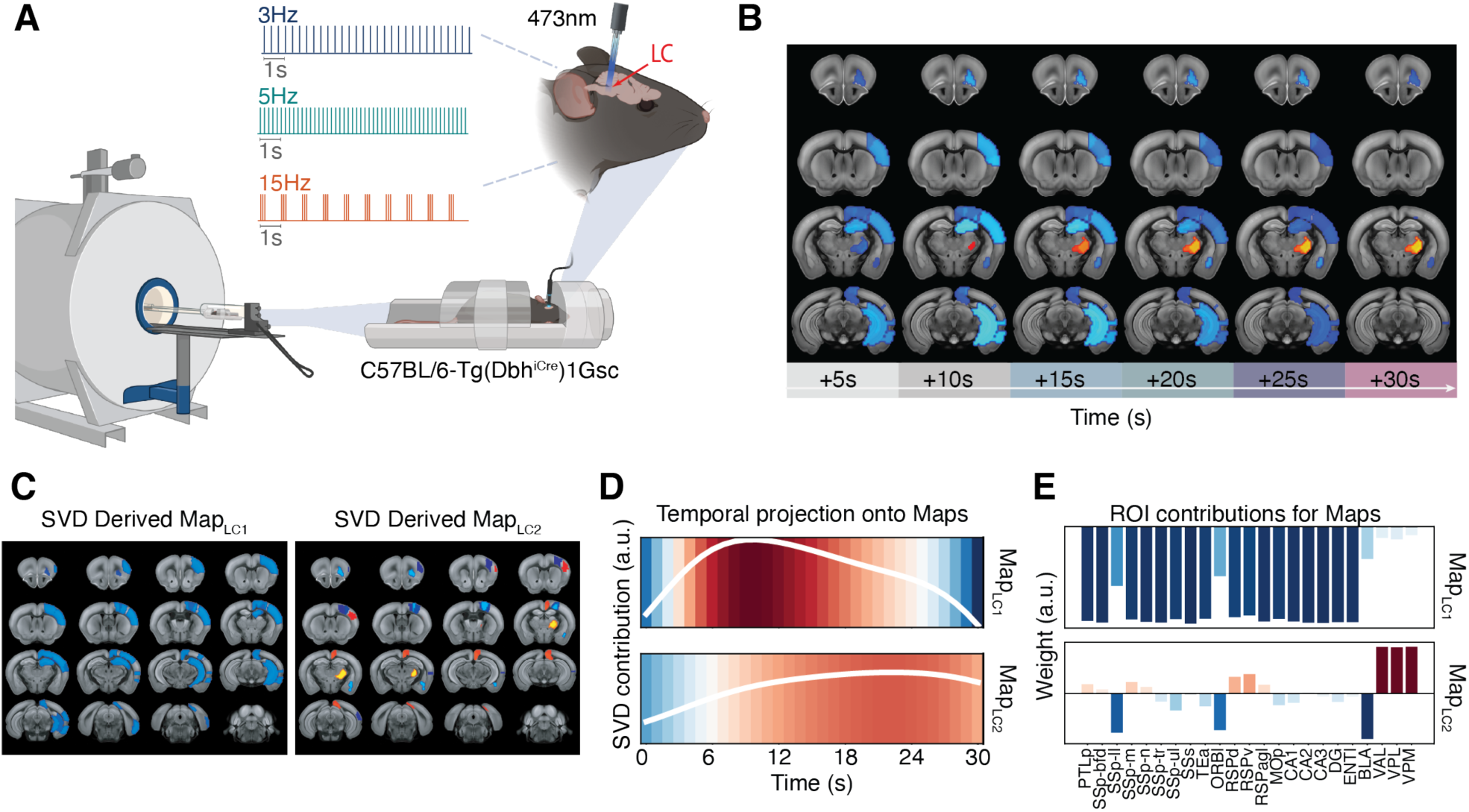
O**p**togenetically**-evoked LC network dynamics reveal a two-component, frequency-invariant spatiotemporal signature**. **A.** Experimental design. Mice expressing channelrhodopsin-2 in LC neurons were stimulated optogenetically at 3, 5, or 15 Hz during simultaneous whole-brain fMRI under light isoflurane anesthesia (9 cycles of 30-s ON/OFF per session). **B.** LC-evoked BOLD responses across stimulation frequencies. Regions consistently modulated relative to sham controls (voxelwise Z > 3.1) exhibited a stereotyped temporal progression from early cortical/hippocampal suppression (blue) to delayed thalamic activation (orange). **C.** Spatial structure of the two dominant SVD components. **D.** Temporal profiles of the SVD components. **E.** ROI-wise spatial weights for the two signature components.

We identified brain regions showing significant LC-evoked modulation relative to sham controls (voxelwise Z > 3.1) across all three stimulation frequencies. The corresponding BOLD time series were z-scored and aligned to stimulation onset (Fig 1B). Regions of interest (ROIs) were retained only if they exhibited high spatial consistency throughout the stimulation period, yielding a set of 23 regions spanning primary and secondary somatosensory cortices, association cortices, hippocampal/limbic structures, and thalamic relay nuclei. To characterize the spatiotemporal structure of the LC response, we applied a singular value decomposition (SVD) to the BOLD activity to the first 30 s after stimulation onset (z-scored per ROI). The number of components was selected to explain 95% of cumulative variance, resulting in two dominant temporal modes that together accounted for 98.7% of the variance. Spatial weights for each mode were obtained by regressing ROI time series onto the corresponding temporal mode, producing one weight per ROI.

This analysis revealed two LC-evoked network components with distinct temporal and spatial structures, hereafter referred to as Map_LC1_ and Map_LC2_. Map_LC1_ captured a negative BOLD deflection in somatosensory and hippocampal regions (primary and secondary somatosensory cortex, CA2/CA3), emerging immediately following LC activation. Map_LC2_ captured a positive BOLD deflection centered on thalamic relay nuclei (VPM, VAL, VPL), which emerged with a temporal delay as the initial cortical suppression dissipated (Fig. 1C-E).

Together, these two SVD-derived maps provide a compact representation of the stereotyped, frequency-invariant LC network response. This low-dimensional template serves as a reference for identifying LC-driven network dynamics across experimental conditions and physiological states.

### 2. The optogenetic LC signature re-emerges during endogenous LC fluctuations

Having established a two-component spatiotemporal LC signature under optogenetic stimulation, we next asked whether the same signature emerges during endogenous fluctuations of LC activity. Endogenous LC surges are prominent during non-rapid-eye-movement (NREM) sleep, during which infraslow modulations of LC firing (in the range of 0.02 Hz) and noradrenaline release generate alternating substates with distinct arousability and electrophysiological profiles [21]. Urethane anesthesia recapitulates these dynamics, producing two alternating brain states with distinct spectral power profiles that reflect cyclic fluctuations in cortical and hippocampal activity [22], [23]. To test whether urethane provides a stable experimental framework for sampling endogenous LC fluctuations, we recorded LC calcium dynamics using fiber photometry of jGCaMP8s [21] in 4 mice, first during natural sleep, then under urethane anesthesia (1.7–1.8 g/kg). As expected, LC calcium surges occurred on an infraslow timescale (∼0.02 Hz) during NREM sleep but were largely absent during REM [21] (Fig. 2A). Under urethane, we observed spontaneous, large-amplitude surges with comparable peak ΔF/F₀ but slower, more variable kinetics (∼8–25 min; Fig. 2B). These surges coincided with a shift from slow oscillation (0.75–1.5 Hz) to theta (4–6 Hz) dominance in cortical EEG and hippocampal LFP (Fig. 2E,F; n = 232 surges), consistent with urethane-associated state transitions [23].

**Figure 2.**
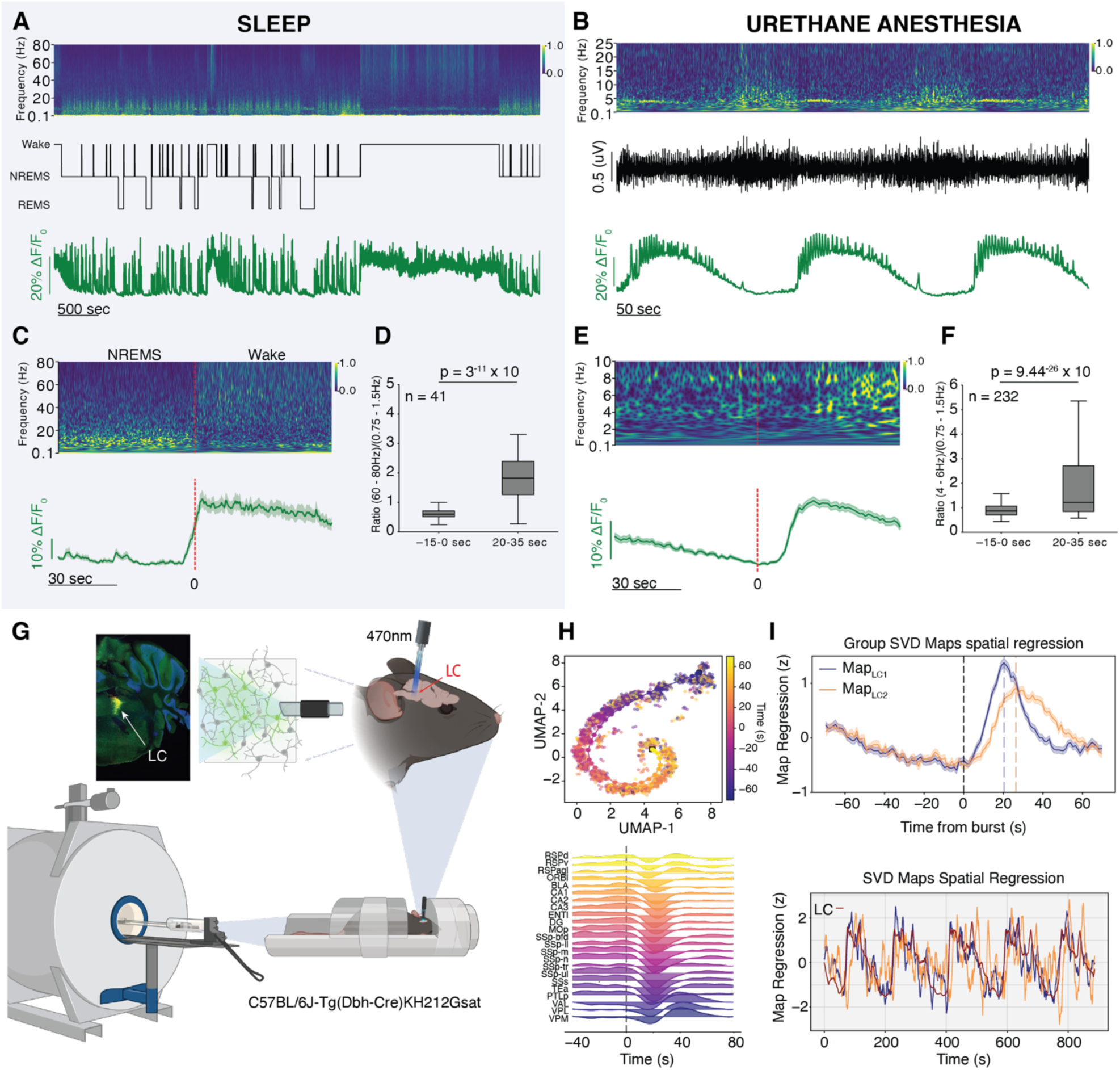
E**n**dogenous **LC-driven dynamics during urethane anesthesia. A.** Representative polysomnographic recordings combined with LC fiber photometry from an animal during an undisturbed sleep–wake cycle. **Top:** Time–frequency spectrogram of the bipolar electroencephalogram (EEG), illustrating state-dependent changes in cortical oscillatory activity. **Middle:** Corresponding hypnogram showing three vigilance states (wake, NREMS, REMS) as determined by polysomnographic recordings. **Bottom:** ΔF/F₀ photometry signal recorded from the LC. **B.** Representative electrophysiological recordings combined with LC fiber photometry of jGCaMP8s from an animal under urethane anesthesia. **Top:** Time–frequency spectrogram of the bipolarized electroencephalogram (EEG), illustrating state-dependent changes in cortical oscillatory activity. **Middle:** Corresponding bipolarized EEG trace. **Bottom:** ΔF/F₀ photometry signal recorded from the LC. **C.** Mean EEG spectrograms and LC ΔF/F₀ signals aligned to n = 41 NREMS-to-wake transitions detected across 9 animals. The dotted red line indicates the time at which the transition was identified based on the hypnogram. **D.** Quantification of EEG spectral dynamics across sleep state transitions, expressed as the power ratio between the gamma (60–80 Hz) and slow oscillation (0.75–1.5 Hz) frequency bands, calculated during the –15 to 0 s and 20 to 35 s time windows relative to the state transition (n = 41 detected NREMS-to-wake transitions). Statistical comparisons were performed using a Wilcoxon signed-rank test. **E**. Mean EEG spectrograms and LC ΔF/F₀ signals aligned to n = 232 surges between the two distinct urethane-induced brain states detected across 4 animals. The dotted red line indicates the time at which the transition was detected based on surge in the LC ΔF/F₀ signal. **F**. Quantification of EEG spectral dynamics across state transitions, expressed as the power ratio between the theta (4–6 Hz) and slow oscillation (0.75–1.5 Hz) frequency bands, calculated during the –15 to 0 s and 20 to 35 s time windows relative to the urethane transition (n = 232 detected surges). Statistical comparisons were performed using a Wilcoxon signed-rank test. **G.** Experimental setup for simultaneous LC fiber photometry (jGCaMP8s) and whole-brain fMRI in urethane-anesthetized mice**. H. Top:** Nonlinear dimensionality reduction (UMAP) applied to whole-brain fMRI trajectories aligned to LC calcium surges (n = 184 events across N = 9 mice). Each point represents a fMRI timepoint (0 to +70 s), colored by time relative to LC surge onset. **Bottom:** ROI-wise fMRI responses (mean ± SEM) for representative cortical, thalamic, hippocampal and amygdalar regions aligned to LC surge onset, illustrating coordinated and temporally ordered propagation of activity across the brain. **I.** Time courses for Map_LC1_ and Map_LC2_ derived by regressing the two optogenetically defined LC SVD components onto the fMRI data. **Top**: Group-averaged similarity traces reveal a robust temporal ordering of the two LC-driven components, with Map_LC1_ emerging earlier and Map_LC2_ following at a delayed peak. **Bottom**: Representative single-animal examples showing stable expression of both components, closely tracking the simultaneously recorded LC calcium signal.

Together, these data demonstrate that urethane preserves robust, state-dependent endogenous LC fluctuations suitable for event-triggered fMRI analysis. We therefore performed simultaneous LC fiber photometry and whole-brain fMRI in urethane-anesthetized mice (N = 9; Fig. 2G) to test whether the optogenetically derived LC signature re-emerges during spontaneous LC activity.

Across all urethane fMRI sessions, we acquired 13 h 15 min of data. Infraslow LC fluctuations were present in the photometry signal for 8 h 30 min in total, during which we detected 184 LC surges using a custom onset-detection algorithm (see Methods). To assess whether LC surges evoke consistent brain-wide reorganization across animals, we aligned whole-brain fMRI volumes to surge onset and embedded the resulting event-centered trajectories using nonlinear dimensionality reduction (UMAP). This unsupervised analysis revealed a reproducible spiral-shaped manifold across all animals, consistent with a stereotyped and cyclic evolution of whole-brain activity following endogenous LC activation (Fig. 2H).

We next tested whether these stereotypical dynamics coincide with the optogenetically derived LC-signatures (Map_LC1_ and Map_LC2_). For each animal, we projected these components onto the fMRI data via spatial regression, yielding scalar similarity time courses that quantify the presence of each LC-driven pattern, following map-projection methods used in prior work [24]. Both Map_LC1_ and Map_LC2_ expression time courses showed robust correlations with the simultaneously recorded LC calcium signal at the individual and group level (Map_LC1_: r = 0.438 ± 0.042, p = 9.35×10^-11^; Map_LC2_: r = 0.571 ± 0.029, p = 1.2 ×10^-17^), indicating that both components of the LC signature are present during endogenous LC fluctuations. When averaging similarity traces time-locked to LC bursts, Map_LC1_ peaked earlier than Map_LC2_, which demonstrates a consistent Map_LC1_→Map_LC2_ ordering during optogenetics and under urethane anesthesia (Fig. 2I).

We next examined the contribution of the global signal (GS) to LC-related large-scale BOLD coupling. To this end, we quantified the variance shared between each spatial map and the GS and reassessed LC–map coupling after controlling for GS effects. Map_LC1_ similarity was strongly dominated by global signal fluctuations (mean R² = 0.996). Nevertheless, its coupling with LC activity remained significant after GS control using both partial correlation and voxelwise GS regression. In contrast, Map_LC2_ showed substantially lower shared variance with the global signal (mean R² = 0.20), and its coupling with LC activity likewise persisted after GS regression. Together, these results indicate that LC–map coupling is not only attributable to global BOLD fluctuations, and reflects structured, map-specific network dynamics.

### 3. LC signature dynamics across naturalistic sleep–wake state transitions in mice

We next asked whether the LC-derived network signature also emerges during spontaneous brain-state transitions. The LC is known to modulate shifts between global states, adjusting arousal levels and coordinating changes in large-scale network configuration [25]. In the sleep–wake cycle, this role is particularly well-defined: electrophysiological and photometric recordings show that brief increases in LC activity reliably precede transitions from NREM sleep to wakefulness or micro-arousals, and that infraslow fluctuations of LC firing modulate the probability of sensory awakening [16], [17], [21], [26]. Because these transitions are accompanied by clear, time-locked changes in LC output (Fig. 2A-D), the sleep–wake cycle provides a suitable model for testing whether the optogenetically derived LC signature also appears during naturally occurring state transitions. To address this question, we analyzed the simultaneous ECoG–fMRI publicly available dataset reported by Yu et al. [27] (N = 21). Animals were head-fixed and extensively habituated to the scanner, resulting in long, stable epochs of wakefulness and NREM sleep, with REM episodes occurring less frequently. For each animal, we quantified LC-signature expression using the same map-projection approach described above. Sleep–wake transitions were extracted from the polysomnographic hypnogram, and map-expression time courses were z-scored, aligned to each transition, and aggregated across animals (n = 1075 NREM→Wake transitions; n = 1089 Wake→NREM transitions; Fig. 3B,C).

**Figure 3.**
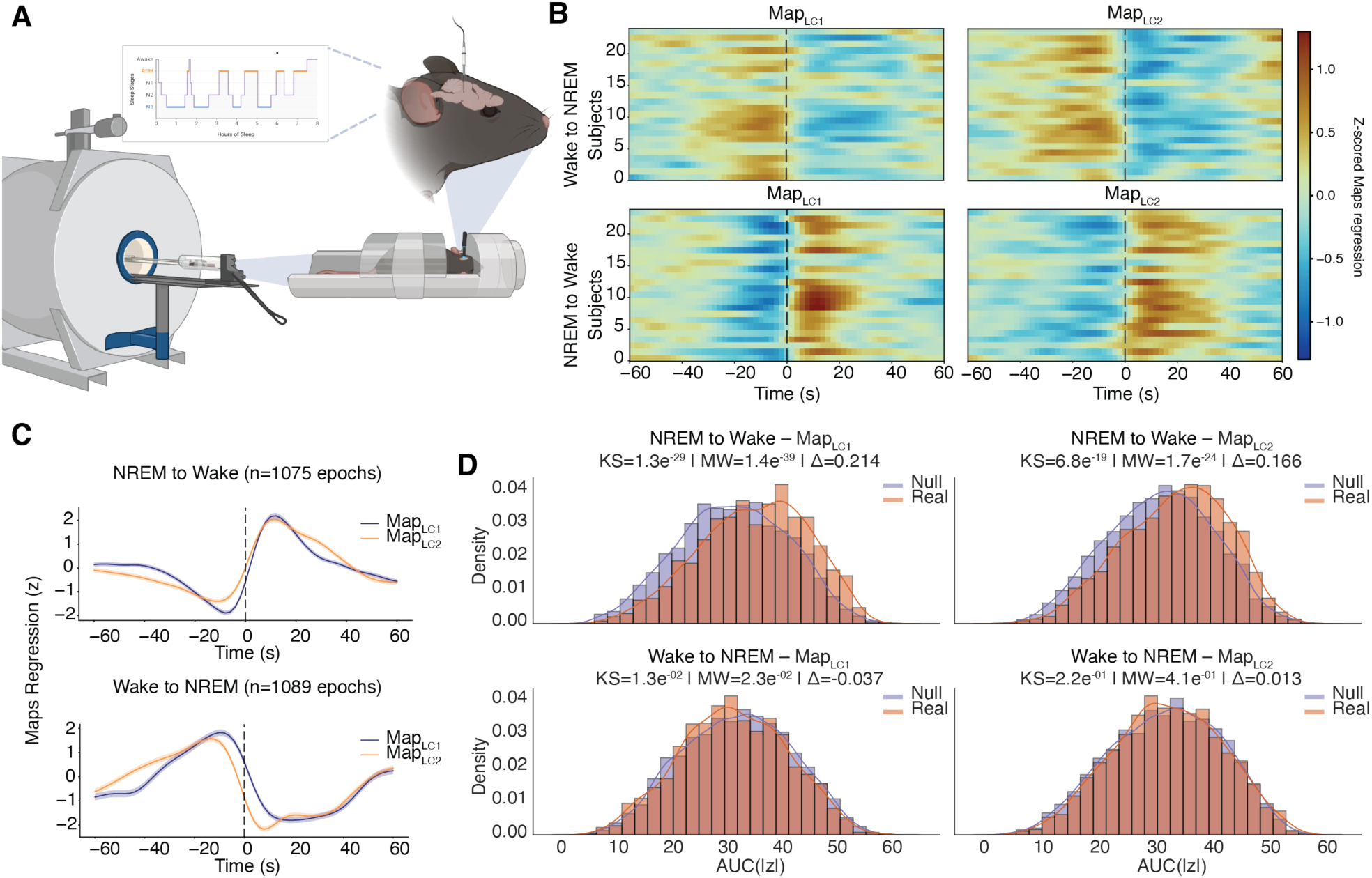
L**C-derived signature is systematically modulated during spontaneous sleep–wake transitions. A.** Schematic representation of the experimental setup for simultaneous ECoG–fMRI recordings in head-fixed, habituated mice. **B.** Event-aligned regressors showing the expression of the two optogenetically derived LC signature components (Map_LC1_, Map_LC2_) projected onto whole-brain fMRI data. Each row corresponds to a transition in an individual animal, aligned to polysomnographically defined Wake→NREM (top) or NREM→Wake (bottom) transitions (time 0). **C.** Averaged LC-map regression time courses across all transitions. NREM→Wake transitions show a rapid and sustained increase in both LC signature components, whereas Wake→NREM transitions display symmetric, prolonged suppression. **D.** Event-wise AUC analysis comparing real transitions (orange) to a randomly sampled null model (purple).

To determine whether LC-signature expression deviated systematically from baseline during these transitions, we quantified event-wise map engagement using a baseline-corrected area-under-the-curve (AUC) metric, defined as the integrated magnitude of map-expression over the post-transition window. Real AUC distributions were compared against a randomly sampled null model to assess whether LC-related network activity was modulated above chance level. This analysis revealed a highly stereotyped and state-dependent modulation of LC-signature expression. During NREM→Wake transitions, both LC signature components showed strong positive deviations relative to the null model. The effect was most prominent for Map_LC1_ (Δ = 0.214; Kolmogorov–Smirnov test, p ≈ 1 × 10^-29^; Mann–Whitney U test, p ≈ 1 × 10^-39^), with Map_LC2_ also showing a robust increase (Δ = 0.166; Kolmogorov–Smirnov test, p ≈ 7 × 10^-19^; Mann–Whitney U test, p ≈ 2 × 10^-24^)(Fig, 3D top panels).

In contrast, during Wake→NREM transitions, map expression showed only weak deviations relative to the null distribution. Map_LC1_ exhibited a small but statistically significant shift (Δ = −0.037; Kolmogorov–Smirnov test, p ≈ 1 × 10^-2^; Mann–Whitney U test, p ≈ 2 × 10^-2^), whereas Map_LC2_ showed a non-significant change (Fig. 3D, bottom panels).

Together, these results demonstrate that the two-component LC signature identified under optogenetic stimulation reappears spontaneously during natural sleep–wake transitions. The polarity and temporal structure of these effects form a strikingly organized pattern: LC-signature expression is suppressed when entering sleep and enhanced during awakening. This organization closely mirrors well-characterized electrophysiological patterns of LC activity, indicating that the fMRI-derived LC signature components capture physiologically meaningful noradrenergic network dynamics during spontaneous brain-state transitions.

### 4. LC signature dynamics across naturalistic sleep–wake state transitions in humans

To determine whether the LC signature generalizes to humans, we applied the same analytical framework to human sleep fMRI (Fig. 4A). We analyzed a publicly available dataset from Horikawa et al. [28] (N = 3 participants, n = 51 scans) that includes whole-brain fMRI with simultaneous polysomnography (PSG).

**Figure 4.**
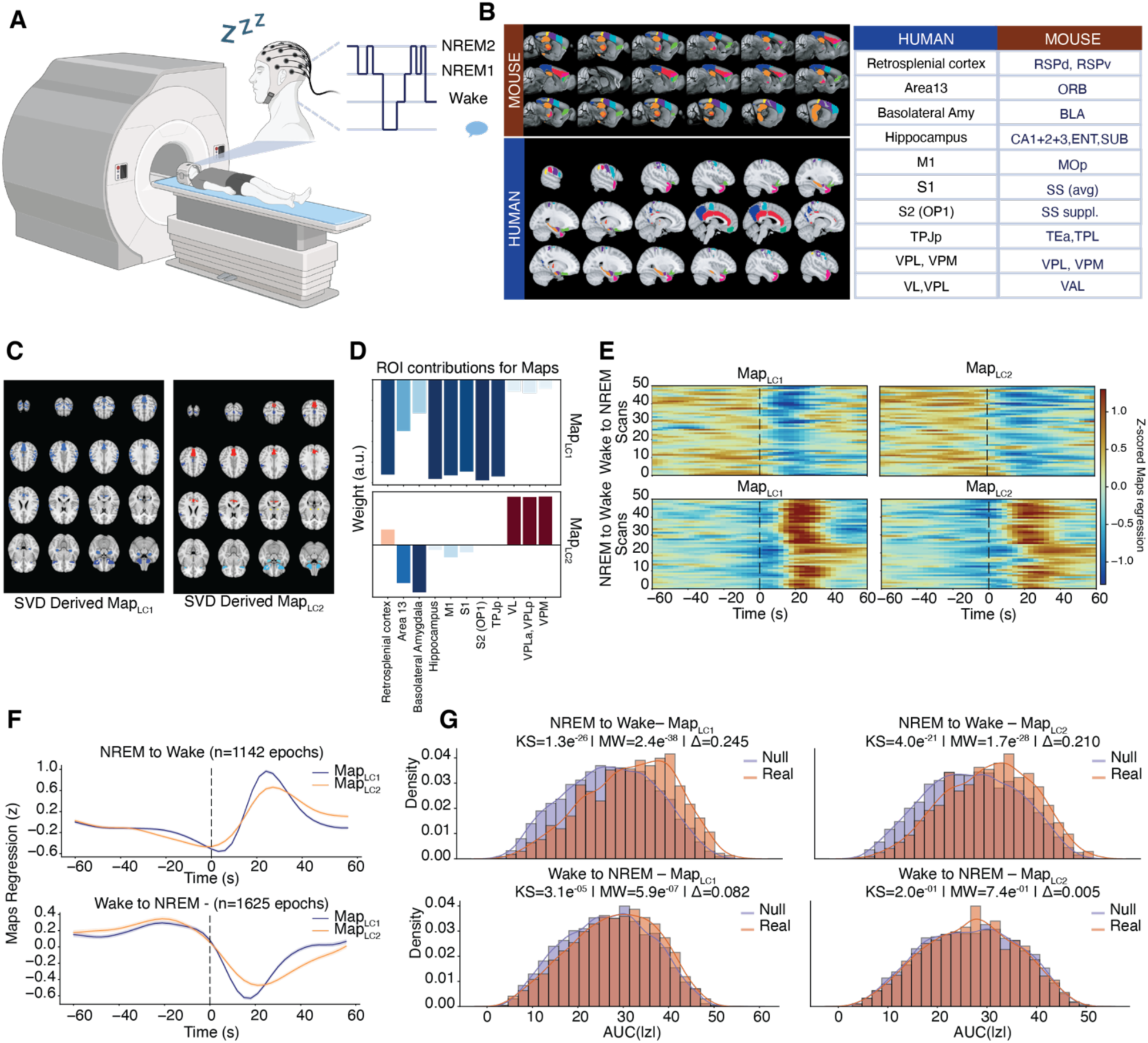
C**r**oss**-species expression of the LC signature during human sleep–wake transitions. A.** Schematic representation of the experimental setup for simultaneous polysomnography (EEG/EMG) and whole-brain fMRI in sleeping human participants. **B. Left**: Cross-species anatomical correspondence between mouse and human regions contributing to the LC signature. **Right**: ROI homology table based on structural and functional correspondence. **C.** Human-space LC signature maps derived by transferring SVD weights from mouse to human homologous ROIs, preserving polarity and magnitude. **D.** ROI-level contributions to human Map_LC1_ and Map_LC2_. **E.** Event-aligned LC-map expression during Wake→NREM (**top**) and NREM→Wake (**bottom**) transitions. Each row represents an individual transition. **F.** Average LC-map regression time courses across all human transitions. **G.** Event-wise AUC comparison of real transitions (orange) against a randomly sampled null model (purple).

To transfer the mouse-derived LC signature to human brain space, we used a cross-species homology table based on Balsters et al. [29], which identifies structurally and functionally corresponding regions between mice and humans. For each mouse ROI, we identified its human anatomical homologue and transferred the SVD-derived weights while preserving their magnitude and polarity for both Map_LC1_ and Map_LC2_. Thalamic relay nuclei with anatomical correspondence were also included to maintain the thalamocortical structure of the LC signature (Fig. 4B-D; see Methods).

The resulting human-space LC signature components were then projected onto each fMRI volume using spatial regression, yielding time courses of LC map-expression across the recording. Map-expression trajectories were z-scored, aligned to PSG-defined transitions, and analyzed using the same event-wise AUC and permutation framework applied to the mouse data.

Consistent with the rodent findings, LC-signature expression exhibited a robust, state-dependent modulation across human sleep–wake transitions (Fig. 4E,F). During NREM→Wake transitions, both LC signature components displayed pronounced positive deviations in AUC relative to their null distributions. Map_LC1_ showed a large right-shifted AUC distribution (Δ = 0.245; Kolmogorov–Smirnov test, p ≈ 1 × 10^-26^; Mann–Whitney U test, p ≈ 2 × 10^-38^), indicating robust reinstatement of LC-network engagement at arousal onset. Map_LC2_ exhibited a similarly strong effect (Δ = 0.210; Kolmogorov–Smirnov test, p ≈ 4 × 10^-21^; Mann–Whitney U test, p ≈ 2 × 10^-28^). During Wake→NREM transitions, AUC values showed much smaller deviations from the null distribution (Map_LC1_: Δ = 0.082; Map_LC2_: Δ = 0.005) (Fig. 4G).

These findings reveal a striking cross-species conservation of LC-driven network dynamics. Human NREM→Wake transitions are characterized by a robust increase in LC-signature expression, whereas Wake→NREM transitions show suppression of the same network. The polarity, timing, and organization of these effects closely mirror the patterns observed in rodents, suggesting that the optogenetically derived LC signature captures an evolutionarily conserved architecture of noradrenergic state modulation across mice and human

#### LC signature dynamics tracks neuromodulatory bursts in awake human fMRI

To test whether the LC-derived signature reflects endogenous noradrenergic fluctuations during wakefulness, we analyzed a 7T resting-state fMRI dataset originally reported by Hearne et al. [30] and reprocessed following the analytical framework of Munn, Müller et al. [25] (N = 59 healthy adults; TR = 586 ms). This dataset includes preprocessed time series from the LC and the cholinergic basal nucleus of Meynert (BNM). Following the latter study, phasic arousal events were identified as rapid, transient increases in the temporal derivative of each neuromodulatory signal. Here, we focused specifically on LC-dominant bursts, defined as events in which LC activity exhibited a prominent phasic increase while BNM activity showed no concomitant increase or was reduced.

Using the event timings for each participant, we projected the two optogenetically derived LC signature components, previously translated into human space using cross-species homology, onto the whole-brain fMRI volumes via spatial regression. This resulted in two similarity time courses per participant, reflecting the moment-to-moment expression of each LC component.

Focusing on LC-dominant bursts, we extracted peri-event segments and computed event-locked averages of LC-signature expression. Both components exhibited clear, time-locked modulations around burst onset, with distinct temporal profiles (Fig. 5B). To assess whether these modulations reflected genuine neuromodulatory engagement rather than nonspecific fMRI fluctuations, we constructed a subject-specific null model consisting of randomly sampled control windows matched in number and duration, following the same AUC framework previously applied to the other datasets.

**Figure 5.**
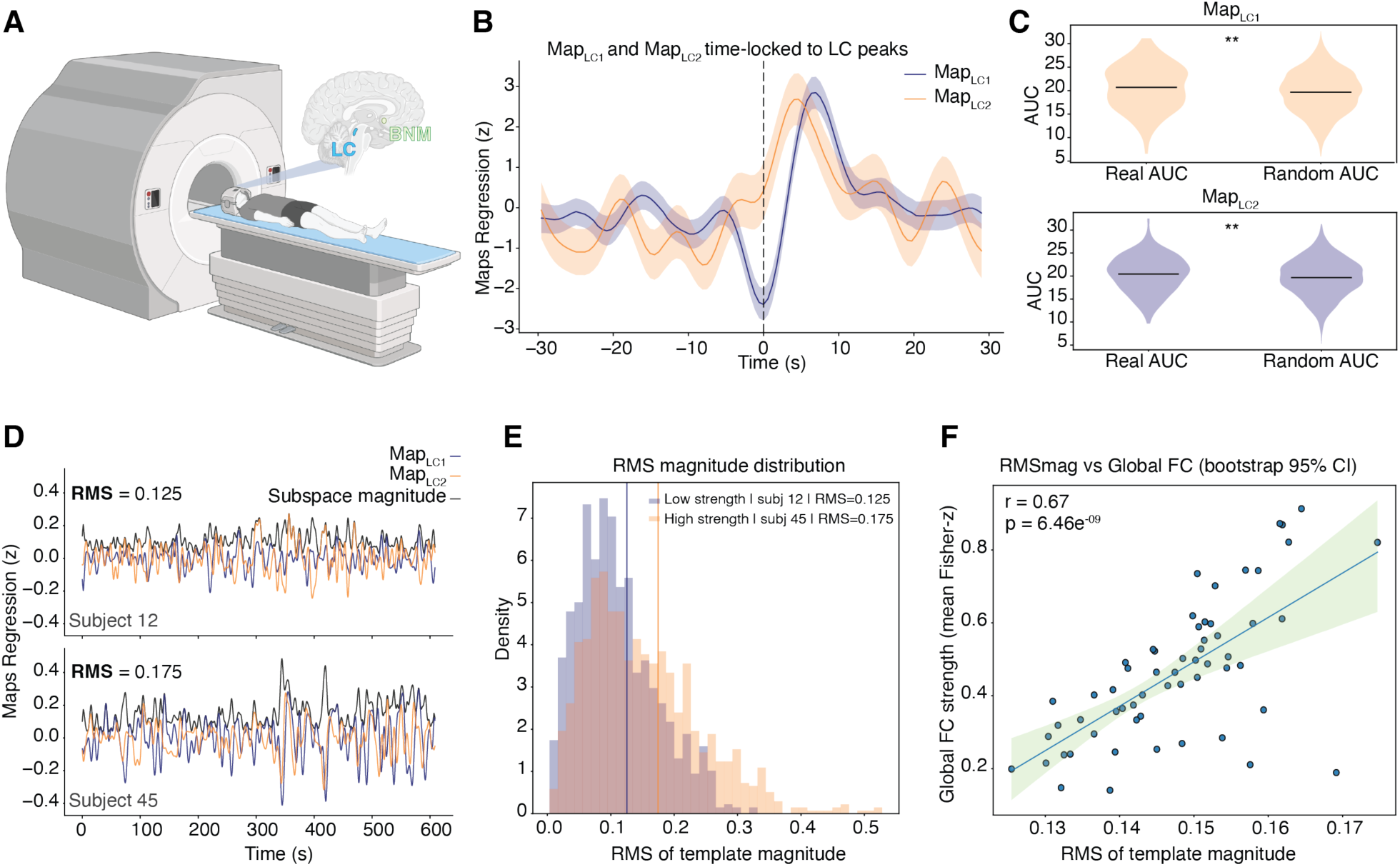
L**C-derived signature tracks spontaneous neuromodulatory bursts and reorganizes large-scale connectivity in awake humans. A.** Schematic of the 7T resting-state fMRI experiment in awake humans. Spontaneous neuromodulatory activity was measured from the locus coeruleus (LC) during rest. **B.** Event-aligned expression of the two LC-derived signature components (Map_LC1_, Map_LC2_) during LC activation events, showing robust modulation of both maps around burst onset (time 0; dashed line). Shaded areas indicate ± SEM. **C.** Baseline-corrected event-wise AUC comparisons between real LC–dominant events and matched random windows. Double asterisks indicate p < 0.01 (Mann–Whitney U test). **D.** Representative example subjects illustrating map expression time series during rest. For each subject, Map_LC1_(blue) and Map_LC2_(orange) expressions are shown together with their combined magnitude (black). Subjects differ in their overall LC state expression strength, summarized as the RMS of this magnitude across time. **E.** Distribution of RMS LC state expression magnitude for two representative subjects with low (blue) and high (orange) expression strength, illustrating inter-individual variability in the typical amplitude of LC-related state expression. Vertical lines indicate subject-specific RMS values. **F.** Relationship between LC state expression strength and global functional connectivity (FC). Each dot represents a participant (N = 59). Global FC strength was computed as the mean Fisher z–transformed Pearson correlation across all ROI pairs. LC state expression strength positively correlated with global FC strength.

Event-wise, baseline-corrected AUC analysis computed over the post-event window revealed selective and map-specific recruitment of the LC signature during phasic noradrenergic events. For LC-dominant bursts, both LC signature components showed significantly greater AUC values compared with randomly sampled control windows (Map_LC1_: Mann–Whitney U test, p ≈ 3 × 10^-4^; Kolmogorov–Smirnov test, p ≈ 3 × 10^-3^; Map_LC2_: Mann–Whitney U test, p ≈ 8 × 10⁻³; Kolmogorov–Smirnov test, p ≈ 1 × 10⁻²), indicating robust engagement of the LC-derived spatial signature during endogenous LC bursts (Fig. 5C).

#### Functional-connectivity consequences of LC-pattern expression in awake humans

Having established that the LC-derived signature tracks spontaneous neuromodulatory bursts in awake humans, we next asked whether the overall expression of this signature during a scan systematically influences commonly used rs-fMRI metrics of large-scale network organization. Because LC-signature expression produces coordinated BOLD fluctuations across distributed regions, greater cumulative expression of this signature during a scan could increase the shared variance between regional time courses and thus contribute to higher functional connectivity estimates. In this framework, functional connectivity (FC) reflects not only stable anatomical and network structure but also the aggregate impact of transient neuromodulatory state shifts that unfold during rest.

To test this idea, we quantified the overall LC state expression strength for each participant, defined as the average magnitude of LC-related network pattern expression over the scan. We then assessed whether inter-individual variation in this measure predicted corresponding differences in whole-brain functional connectivity. Specifically, we projected z-scored ROI activity onto the two LC-derived maps, yielding two map-expression time series, and summarized each participant’s expression by computing the RMS of 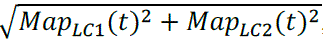, which captures the overall amplitude of LC-related network engagement independent of sign or temporal ordering. In parallel, we computed a global FC strength measure for each participant as the mean Fisher z–transformed Pearson correlation across all cortical ROI pairs (Fig. 5D,E).

We observed a significant positive correlation between LC subspace strength and cortical FC strength (r = 0.67, p = 6.46×10^-9^, N = 59). Participants exhibiting stronger LC-related state expression also showed higher mean functional connectivity across the scan, consistent with the notion that phasic LC dynamics promote network integration (Fig. 5F).

Together, these findings suggest that endogenous neuromodulatory dynamics shape fluctuations in large-scale connectivity during rest. Importantly, this relationship highlights a critical consideration for interpreting rs-fMRI: variability often attributed to stable individual differences may, in part, reflect differences in internal state expression and neuromodulatory tone. In the context of emerging individualized fMRI-based biomarkers, such endogenous state fluctuations represent a biologically meaningful source of variance that must be accounted for.

## Discussion

In this work, we show that LC activity engages a conserved, low-dimensional signature that recurs across experimental manipulations, physiological states, and species. By defining this signature under optogenetic control and tracking its expression during endogenous LC fluctuations, natural sleep–wake transitions, and awake human rest, we demonstrate that noradrenergic activity leaves a structured, temporally ordered imprint on whole-brain fMRI dynamics, indicating that neuromodulatory state fluctuations constitute a fundamental, yet often unaccounted, source of variance in resting-state connectomics [1], [24].

### LC activation engages a conserved two-component network with state-dependent temporal dynamics

Optogenetic activation of the LC, as well as endogenous LC fluctuations, consistently engaged two large-scale LC signature components in the fMRI signal. One component was characterized by a suppression of activity in cortical and hippocampal regions, while the other involved increased activity in thalamic nuclei. Similar spatial patterns have been described in previous fMRI studies of natural arousal state transitions [24], [31]. Here, we show that these components are not restricted to a single behavioral or physiological condition, but recur across optogenetic stimulation, spontaneous LC activity, sleep–wake transitions, and awake human rest. Although the spatial organization of these components remained stable across conditions, their temporal relationship depended on physiological state. In awake humans, LC-related fMRI responses followed the sequence described in arousal studies, with thalamic activation preceding cortical suppression [31]. In sleep, the two components unfolded with largely overlapping time courses. Under anesthesia, however, the temporal order was reversed, with suppression of cortical and hippocampal activity preceding thalamic activation. Importantly, this pattern was observed under both isoflurane and urethane anesthesia, despite their distinct molecular mechanisms. This consistency points to a broader reorganization of thalamo-cortical dynamics during anesthesia, rather than a drug-specific effect, in line with previous work describing altered thalamocortical communication during anesthetic-induced loss of consciousness [32], [33].

### Urethane anesthesia preserves sleep-like LC dynamics and associated network reorganization

We found that large and slow LC activity fluctuations appeared in urethane-anesthetized animals. This provided an unprecedented opportunity to examine the fMRI signature of fluctuating brain states induced by endogenous LC activity patterns. Endogenous, infraslow LC fluctuations have so far been described for the polysomnographically defined state of NREM sleep, during which they induce alternating substates that differ in arousability [34], [35], autonomic control [35], [36], REM sleep propensity [21] and glymphatic clearance [37]. These features have been proposed to partition NREM sleep into alternating introspective and engaged states, yet this has so far not been accessible to investigation via fMRI. Urethane anesthesia has been used as a pharmacological model for cyclical NREM and REM sleep-like states [23]. The emergence of LC-dependent brain state fluctuations under this condition provides a valuable experimental analogue to probe the dynamics of sleep-like substates, for which we offer here several noteworthy findings. First, urethane anesthesia-associated brain state dynamics indeed overlap with the ones of NREM sleep, suggesting that they represent a valuable platform to probe LC-driven modulation of global brain activity. Second, this model will allow to study LC’s influence on arousal-related network dynamics in the absence of confounding sensory or motor attributes of natural NREM sleep, possibly linking circuit physiology of arousal to whole-brain imaging. Third, our data clarify that the theta-power-enriched urethane state is accompanied by elevated LC activity, which contrasts with the silence of LC activity known for natural REM sleep [21], [38]. These finding challenges long-held propositions that urethane induces REM sleep-like states and rather qualifies the theta-enriched state as an activated urethane substate with possibly higher arousability. This interpretation is in line with findings showing that LC stimulation during isoflurane anesthesia drives the emergence of theta rhythms and accelerates recovery [39].

### Cross-species conservation of LC-mediated network states

Most comparative studies of mouse and human functional connectivity have focused on describing similarities in large-scale network organization and resting-state architecture across species [40], [41]. Here, we took a different approach by starting from a network pattern defined under causal LC control in mice and asking whether this same pattern could be identified in specific time-locked events in human fMRI data. To this end, we projected the optogenetically derived LC signature components from mouse brain space onto human recordings, and we found that the LC signature was similarly modulated during a comparable, temporally aligned brain-state transition, namely NREM sleep→Wake transition. Interestingly, in mice as well as in humans, transitions from wakefulness to NREM sleep were associated with a reduced presence of the same pattern. The polarity and temporal organization of these effects closely mirror well-established electrophysiological observations showing elevated LC activity during wakefulness and reduced firing during NREM sleep in mammals [15], [16], [17], [36]. The presence of this correspondence across species suggests that the LC-related network dynamics captured by the template reflect a conserved aspect of noradrenergic state regulation, rather than species-specific features or experimental artifacts. At the same time, several limitations should be considered when interpreting these cross-species results. Anatomical homology tables provide a practical framework for mapping networks between mouse and human brain space, but they inevitably rely on approximations [42]. Moreover, differences in LC physiology across species, including firing patterns, baseline activity levels, and adrenergic receptor expression, are likely to influence how LC activity is reflected in the fMRI signal [12], [43], [44]. Methodological differences between mouse and human imaging, such as magnetic field strength, spatial resolution, and acquisition protocols, may further contribute to quantitative differences in the magnitude or timing of LC-signature expression. These factors are expected to affect the expression of the signal without altering the overall pattern of modulation observed across species.

### Partial overlap between LC signatures and global signal reveals distinct arousal-related network components

Growing evidence suggests that the global fMRI signal contains biologically meaningful components linked to arousal, in addition to systemic physiological contributions. Simultaneous EEG–fMRI studies have shown that global BOLD fluctuations covary with vigilance and arousal state [24]. Similarly, pharmacological and behavioral manipulations of arousal modulate both global signal amplitude and large-scale connectivity patterns [1], [45]. More recent work further supports the view that global fMRI fluctuations reflect organism-wide arousal processes rather than noise alone [46], [47].

In our data, the network patterns associated with LC activity shared a portion of their variance with the fMRI global signal. Critically, however, LC-related fMRI effects remained significantly correlated with fiber photometry-measured LC activity even after global signal regression. This indicates that global signal regression does not fully eliminate LC-driven effects but rather removes only the shared variance between arousal-related LC dynamics and global fluctuations. Importantly, the two LC-derived maps differed in how strongly they overlapped with the global signal, pointing to network-specific noradrenergic effects rather than a single global modulation.

Our findings therefore suggest that LC-driven network dynamics represent a physiologically meaningful component of rs-fMRI that is only partly captured by the global signal. This has important implications: while global signal regression is often assumed to remove arousal-related confounds entirely, our results demonstrate that LC-related arousal signals persist after this preprocessing step. Consequently, global signal regression may inadvertently create a false sense of arousal-related noise removal while leaving substantial LC-driven variance in the data, a confound that could systematically bias functional connectivity interpretations [48], [49].

### Inter-individual LC-state differences predict functional connectivity strength in awake human rs-fMRI scans

In the awake human 7T resting-state dataset, inter-individual differences in LC-state expression during rest were strongly associated with mean functional connectivity strength, indicating that ongoing neuromodulatory state exerts a measurable influence on large-scale functional coupling. Similar state-dependent effects have been reported in previous studies linking arousal and noradrenergic tone to global network organization [24], [50]. These observations support the view that resting-state connectivity reflects not only stable anatomical architecture or long-term synaptic coupling, but also transient fluctuations in neuromodulatory state that unfold during the scan. Variability in functional connectivity across individuals has long been hypothesized to arise from a combination of structural differences and state-dependent factors such as spontaneous arousal fluctuations or differences in vigilance regulation [51], [52]. Our results provide direct empirical evidence for this framework by linking individual differences in functional connectivity to measurable variations in LC-driven state expression. This distinction is particularly relevant in clinical populations, where regulation of arousal via the LC–noradrenergic system is frequently altered. A range of psychiatric conditions have been associated with atypical LC dynamics, including changes in tonic–phasic firing balance and noradrenergic reactivity [43], [44]. Such alterations are likely to bias the prevalence of arousal-related network states during scanning, thereby influencing functional connectivity estimates and complicating the interpretation of group differences.

In this context, LC-derived network signatures provide a quantitative tool to estimate neuromodulatory state contributions to functional connectivity at the level of individual scans. Explicitly accounting for such state-dependent effects may therefore improve the interpretability of resting-state connectivity measures and refine the use of fMRI biomarkers in both clinical and translational research.

## Materials and Methods

### 1. Animals and experimental cohorts

For experiments combining fiber photometry with fMRI, we used mice from the C57BL/6-Tg(Dbh-iCre)1Gsc line provided by Prof. Tommaso Patriarchi (University of Zurich) and maintained on a C57BL/6J genetic background. Mice were initially housed in a temperature and humidity-controlled animal facility under a 12-h light/dark cycle, with lights on at 9:00 a.m. (ZT0), and with food and water available ad libitum. For viral injections and fiber implantations, male and female mice aged 5-7 weeks were transferred to a P2 biosafety room, where they were housed from one day before until three days after surgery. Following surgical procedures, animals were allowed to recover for at least one week. After recovery, mice were transferred to the animal facility at the Center for Biomedical Imaging (CIBM), EPFL (Lausanne), where they were maintained under the same housing conditions (12-h light/dark cycle, ad libitum access to food and water) until the fMRI–fiber photometry experiments. A total of nine mice were included in the in-house fMRI–fiber photometry experiments. In addition, previously published data from mice of the B6.FVB(Cg)-Tg(Dbh-Cre)KH212Gsat/Mmucd line were reanalyzed from Osorio-Forero et al. [21] (Fig. 2A,C,D).

All experimental procedures complied with Swiss National guidelines for animal research and were approved by the Swiss Cantonal Veterinary Office Committee for Animal Experimentation.

### 2. Viral vectors and surgical procedures

Surgical procedures for viral delivery, LC fiber implantation, and electrode implantation were performed following protocols comparable to those previously reported in [21]. Briefly, Dbh-iCre mice received injections of an adeno-associated viral vector (AAV5-hSyn1-dlox-jGCaMP8s-dlox-WPRE-SV40p(A); titer 5.8 × 10¹² viral genomes/mL; total volume 600 nL) into the right LC. Injections were carried out using a fine glass micropipette at the following stereotaxic coordinates: lateral (L) 1.05 mm, anterior–posterior (AP) −5.4 mm, and dorsal–ventral (DV) −3.2 to −2.2 mm, delivered in 0.2 mm steps with 100 nL injected at each depth. After completion of the injection, the pipette was left in place for 10 min to allow diffusion of the virus. Optical fiber implantation was then performed by placing a fiber stub coupled to a cannula (Doric Lenses, MFC_400/430-0.66_3.5mm_ZF1.25(G)_FLT) above the right LC at coordinates L 0.9 mm, AP −5.4 mm, and DV

−2.7 mm. The fiber was lowered at a rate of 1 mm/min using a Kopf stereotaxic apparatus. The implant was secured to the skull with adhesive and embedded in dental cement to ensure long-term stability. For animals that underwent polysomnographic recordings, electrode implantation procedures were identical to those previously described in [21].

### 3. Urethane anesthesia - preparation and injections

For recordings performed under urethane anesthesia, mice received an intraperitoneal (i.p.) injection of a urethane solution (10% w/v in saline; Sigma-Aldrich No. 94300) at a final dose of 1.7–1.8 g/kg. After the animals reached a stable level of anesthesia, body temperature was continuously maintained at 37 °C using a feedback-controlled heating system throughout the recording session.

### 4. Data acquisition for simultaneous fMRI–fiber photometry under urethane (in-house)

#### 4.1 Animal preparation

For fMRI acquisitions, mice were first anesthetized in an induction chamber with 4% isoflurane in a 1:4 O₂-to-air mixture for approximately 4 minutes. Following induction, urethane anesthesia was administered (see ‘Urethane anesthesia - preparation and injections’), after which isoflurane was discontinued. Mice were allowed sufficient time for the urethane anesthesia to stabilize and were then positioned on an MRI-compatible cradle and secured using ear bars to minimize head motion. An MRI-compatible fiberoptic connector was used to couple the patch cord to the implanted optical fiber for simultaneous photometry recordings. Body temperature was maintained at 37 °C throughout the entire scanning session using a circulating warm-waterbed and continuously monitored using a rectal thermometer probe. Respiratory rate was recorded using a pneumatic pressure sensor positioned under the abdomen. After acquisition of all functional and anatomical MRI sequences, animals were transcardially perfused with phosphate-buffered saline (PBS 1M; 20 ml), followed by fixation (4% Paraformaldehyde) when required for histological verification of fiber placement.

#### 4.2 Data acquisition

fMRI data were acquired on a 9.4 T horizontal bore magnet (Magnex Scientific, Yarnton, UK) equipped with a shielded gradient system providing a maximum gradient strength of 660 mT m⁻¹ and a slew rate of 4570 T m⁻¹ s⁻¹ (Bruker B-GA12S HP), interfaced to a Bruker BioSpec console running ParaVision 360 (v3.5). A custom-built, transmit-receive saddle coil was used (JD Coils). The coil is equipped with non-magnetic, variable capacitors to ensure sample-specific impedance matching at the resolance frequency and specifically designed to accommodate the optical fiber and patch cord for simultaneous fiber photometry recordings. After standard scanner adjustments, including frequency calibration and shimming, functional images were acquired using a gradient-echo echo-planar imaging (GE-EPI) sequence. Imaging parameters were as follows: repetition time (TR) = 1200 ms, echo time (TE) = 13 ms, matrix size = 76 × 60, field of view = 19 × 15 mm², resulting in an in-plane resolution of 0.25 × 0.25 mm². Fifteen axial slices were acquired with a slice thickness of 0.5 mm and an inter-slice gap of 0.1 mm, yielding a slice distance of 0.6 mm. Slice acquisition was interlaced, with a single shot per volume and no parallel imaging or multiband acceleration applied. Each functional run consisted of 740 volumes, corresponding to a total scan duration of 15 min. This configuration enabled stable whole-brain functional imaging while maintaining compatibility with simultaneous fiber photometry acquisition.

#### 4.3 Fiber photometry recordings

Fiber photometry recordings were performed using a Doric fluorescence MiniCube system (ilFMC4-G2_IE(400–410)_E(460–490)_F(500–550)_S, Doric Lenses) to monitor low-frequency fluorescence fluctuations during urethane anesthesia. Two signal generators independently modulated the violet (405 nm) and blue (465 nm) excitation LEDs at 211 Hz and 319 Hz, respectively, allowing synchronous demodulation of the isosbestic and activity-dependent channels. The combined modulated light was delivered through a 6.3 m low-autofluorescence mono fiberoptic patchcord (400 µm core, NA = 0.57; Doric Lenses, model MFP_400/430/1100-0.57_6.3m_FC-ZF1.25_LAF) routed inside the MRI scanner and coupled to the implanted optic cannula on the animal’s head. Fluorescence emitted by the sensor was collected through the same fiber and converted into a current signal by the photodetector integrated in the MiniCube. Excitation power at the fiber tip was maintained between 20–30 µW for both wavelengths. Continuous photometry acquisitions ranged from 2 to 3 hours, spanning entire fMRI sessions. In parallel, an external trigger signal marking the onset of the fMRI acquisition was recorded via a separate analog input channel in the Doric console, to enable precise temporal alignment between photometry and fMRI data.

#### 4.4 Histology and immunohistochemistry (in-house)

Following transcardial perfusion, brains were extracted and post-fixed for 24 h in 4% paraformaldehyde (PFA) at 4 °C. Tissue was then cryoprotected in 30% sucrose for 1–2 days and sectioned at 50 µm thickness using a manually guided freezing microtome (Microm). Brainstem sections were collected for subsequent analyses. To assess the colocalization of jGCaMP8s expression with tyrosine hydroxylase (TH) in LC neurons, 50-µm-thick coronal sections corresponding to approximately −5.3 mm from bregma were processed for immunohistochemistry. Endogenous jGCaMP8s fluorescence was enhanced by GFP signal amplification. Sections were washed three times in PBS containing 0.3% Triton X-100, followed by incubation in blocking solution (PBS, 0.3% Triton X-100, 2% normal goat serum) for 1 h. Primary antibodies were then applied overnight at 4 °C under gentle agitation: rabbit anti-TH (1:5,000; Merck Millipore, AB152) and chicken anti-GFP (1:2,000; Abcam, AB13970). After at least 12 h of incubation, sections were washed three times in PBS containing 0.3% Triton X-100 and incubated with secondary antibodies for 1–1.5 h at room temperature on a shaking platform. Secondary antibodies included donkey anti-rabbit Alexa Fluor 647 (1:300; Thermo Fisher Scientific, A31573) and donkey anti-chicken (1:500; Jackson ImmunoResearch, 703-545-155), both diluted in PBS with 0.3% Triton X-100. Sections were subsequently rinsed in 0.1 M PBS and mounted with Mowiol. Fluorescent images were acquired using a confocal microscope (Leica Stellaris 8) through green and far-red emission channels, using LAS X software (v4.6.1.27508).

### 5. Sleep and urethane electrophysiology/photometry recordings (previously published cohort)

Sleep–wake electrophysiology and LC fiber photometry recordings were obtained from the dataset originally reported by Osorio-Forero et al. [21]. All experimental procedures, including data acquisition, sleep staging, and recording hardware, were performed as described in the original study. Among these animals, four had undergone simultaneous electrophysiological recordings and LC fiber photometry under urethane anesthesia. These urethane recordings were not analyzed in the original publication and are analyzed here for the first time. For urethane sessions, recordings were limited to a maximum duration of 5 h after injection. Further details regarding the sleep–wake recordings of this cohort are available in [21].

### 6. Data analysis

#### 6.1 Polysomnography and spectral analysis (previously published cohort)

Polysomnographic scoring procedures were performed as previously described in [21]. To quantify changes in cortical oscillatory activity associated with brain-state transitions under urethane anesthesia or from NREMS to wakefulness, the bipolar EEG signal was converted into time–frequency spectrograms using wavelet transforms based on Gabor–Morlet kernels. Four-cycle wavelets were applied, with Gaussian envelope standard deviations corresponding to frequency resolutions of 0.05 Hz and 0.25 Hz, respectively. Spectral power ratios were then calculated for specific frequency bands: theta (4–6 Hz) relative to slow oscillations (0.75–1.5 Hz) for urethane state transitions, and 60–80 Hz relative to 0.75–1 Hz for NREMS-to-wake transitions. These ratios were computed and compared between two-time windows (−15 to 0 s and 20 to 35 s) aligned to the detected LC surge onset for the urethane data and to NREMS-to-wake transition for the undisturbed sleep data.

All statistical analyses were conducted in Python. Data distribution normality was evaluated using the Shapiro–Wilk test, and variance homogeneity was assessed with Levene’s test. When assumptions of normality were not met, nonparametric Wilcoxon tests were applied.

#### 6.2 Fiber photometry processing

Fiber photometry recordings were preprocessed using a custom pipeline developed in Python to correct slow drifts, normalize fluorescence signals, and align the photometry trace to the corresponding fMRI acquisition. To correct for photobleaching and isolate activity-related fluorescence changes, the activity-dependent channel was fitted to the isosbestic control signal using a second-degree polynomial function, and ΔF/F_0_ was computed as 100 × (F(t) − F^(t)) / F^(t), where F(t) is the recorded fluorescence and F^(t) is the fitted control-derived baseline. Because the experiment focused on very slow fluorescence fluctuations characteristic of urethane anesthesia, the ΔF/F_0_ signal was smoothed using a moving-average filter (150-point window) to suppress high-frequency noise while preserving slow dynamics. The resulting ΔF/F_0_ trace was then z-scored across the recording. Because each fMRI scan lasted 15 min, an external trigger channel was used to identify the exact onset of each MRI acquisition, and the photometry signal was segmented accordingly so that only the portion temporally matched to the fMRI run was retained for further analysis and interpolated to the fMRI repetition time (TR = 1.2 s). LC surge onsets were then identified from the TR-locked photometry signal using a custom detection algorithm designed to capture the beginning of slow LC waves rather than their peak amplitudes. The LC trace was low pass filtered at 0.02 Hz to isolate infra-slow fluctuations, and second derivatives were computed to identify inflection points corresponding to rapid upward transitions in the signal. Candidate onsets were retained only if they were followed by a sustained fluorescence increase of at least 0.6 ΔF/F_0_ within the subsequent 20 fMRI volumes and exceeded the mean baseline level, yielding a set of LC surge onsets temporally matched to individual fMRI volumes for the other analyses. In total, 184 LC surges were detected across all animals.

#### 6.3 fMRI preprocessing and spatial regression (all datasets)

##### 6.3.1 Opto-fMRI Dataset Processing

For the optogenetic dataset, we re-used the publicly available fMRI data from Grimm et al. [18], which had already been preprocessed and co-registered to a common anatomical template. Functional scans were acquired using a gradient-echo EPI sequence (GE-EPI; TR = 1 s, TE = 15 ms, in-plane resolution = 0.22 × 0.20 mm², 20 slices, slice thickness = 0.4 mm, slice gap = 0.1 mm). For each stimulation frequency (3, 5, and 15 Hz), ROI-wise time courses were extracted from the parcellated data and z-scored within each run. Stimulus-locked responses were computed by extracting for each stimulation a 60 s window (TR = 1 s) starting at stimulation onset and averaging across stimulations and runs to obtain a single mean response time course per ROI; this window was chosen to capture the full stimulation cycle in the original design (30 s stimulation ON followed by 30 s OFF). To identify a robust optogenetic response signature that generalized across stimulation frequencies, we quantified ROI-wise similarity across the 3 Hz, 5 Hz, and 15 Hz mean response matrices and retained a “core” set of ROIs showing consistent temporal profiles across datasets (pairwise ROI-to-ROI diagonal correlations; inclusion threshold r > 0.75). The core ROIs were used to define a frequency-invariant response template computed as the average of the three frequency-specific mean responses. To characterize the temporal evolution of this template, the ROI×time matrix (first 30 s post-stimulation) was z-scored per ROI and decomposed using singular value decomposition (SVD). The number of components was chosen to explain 95% of cumulative variance, yielding two dominant temporal modes explaining 98.7% of the total variance. Spatial weights (Map_LC1_ and Map_LC2_) were obtained by regressing each ROI time series onto the corresponding temporal mode, yielding one weight per ROI.

##### 6.3.2 fMRI – Photometry under urethane anesthesia dataset processing and spatial regression

For the in-house combined fMRI–photometry dataset acquired under urethane anesthesia, fMRI preprocessing was performed using FSL (www.fmrib.ox.ac.uk/fsl). Functional images were corrected for head motion using MCFLIRT, spatially smoothed with a Gaussian kernel (FWHM = 0.5 mm), and temporally high-pass filtered with a cutoff of 100 s to remove slow drifts. Independent component analysis was then applied separately to each run, extracting 30 components per scan. Components reflecting motion, physiological noise, or scanner-related artifacts were manually identified and regressed out from the data, together with residual motion-related variance, using linear regression. The resulting functional scans were then registered to the Allen mouse brain template at 200 µm isotropic resolution using FLIRT with 12 degrees of freedom. Fiber photometry data were temporally aligned to each fMRI scan as described above, yielding a TR-locked LC fluorescence trace for each run. Using the detected LC surge onsets, fMRI volumes corresponding to the onset and progression of spontaneous LC activity were identified. These events were used to examine the expression of large-scale brain patterns previously derived from the optogenetic dataset. Specifically, spatial regression was performed between the fMRI data and the SVD-derived spatial maps obtained from optogenetic LC stimulation and registered to the same Allen template space. At each time point, fMRI signals were z-scored across time within each voxel, and the normalized dot product between the map weights and the instantaneous activity vector was computed. This yielded a single similarity value per volume, providing a continuous measure of LC-related pattern expression while minimizing the influence of global amplitude differences and following established map-projection approaches used to track state-dependent network motifs in fMRI.

Similarity time courses were compared to the simultaneously acquired LC photometry signal using zero-lag Pearson correlation. Prior to correlation, both signals were smoothed with an 8 s moving-average window to suppress high-frequency fluctuations not relevant under urethane anesthesia. Statistical significance was assessed using a permutation-based approach: surrogate LC time series were generated by phase randomization (1,000 iterations), preserving the amplitude spectrum while disrupting temporal alignment, and empirical p-values were computed based on the resulting null distribution of correlation coefficients. Group-level inference was performed by Fisher z-transforming run-level correlation coefficients and testing them against zero using a one-sample t-test. Finally, similarity time courses were aligned to LC surge onsets (±70 s) to compute group-average burst-triggered responses for each spatial map, allowing characterization of the temporal evolution of map-expression around spontaneous LC activation events.

##### 6.3.3 fMRI – Sleep Ecog Dataset Processing and Spatial Regression

For the sleep dataset, we re-used publicly available fMRI–ECoG recordings previously reported by the original authors (Yu et al., [27]), which had already undergone preprocessing and sleep-state annotation as described in detail in the original publication. Functional scans were acquired using single-shot echo-planar imaging (EPI; TR = 2 s, TE = 14 ms, flip angle = 70°, matrix size = 90 × 45, nominal in-plane resolution = 200 × 200 μm², 22 slices, slice thickness = 400 μm). Each recording consisted of a single continuous acquisition of 7,200 volumes (4 h). Slice-level trigger pulses were generated by the MRI console and recorded together with electrophysiological signals, enabling precise temporal alignment between fMRI data, ECoG recordings, and sleep scoring. Sleep states (wake, NREM, and REM) and electrophysiological features were derived by the original authors and directly used in the present study. No additional preprocessing of the raw electrophysiological or sleep-scoring data was performed prior to the analyses described below. Because individual animals were not coregistered across subjects, SVD-derived spatial maps obtained from optogenetic LC stimulation were registered separately to each subject’s native fMRI space. Spatial regression was then performed using the same framework adopted for the urethane dataset. Specifically, each voxel time series was z-scored across time, and for each SVD-derived spatial map, a map-specific regression time course was computed as the normalized dot product between the voxel values of the map and the corresponding voxel values of each fMRI volume. This yielded one regression coefficient per volume, quantifying the instantaneous expression of each spatial pattern over time. Transition events (wake↔NREM) were detected at state boundaries and retained only when (i) a full ±60 s window around the transition was available and (ii) the post-transition state persisted for at least 20 s. For each transition, similarity epochs (±60 s) were extracted for all maps and aggregated across subjects. Event-wise effects were quantified using an AUC metric computed on baseline-corrected similarity traces: each epoch was baseline-referenced (−20 to −10 s), z-normalized within the event, and the AUC of |z| was integrated over 0–30 s post-transition. Null distributions were generated by sampling time windows of identical length from the continuous similarity time series, and real versus null AUC distributions were compared using nonparametric tests (Kolmogorov–Smirnov and Mann–Whitney U), complemented by Cliff’s delta as an effect size.

##### 6.3.4 Mouse to Human analogy

To enable direct comparison between mouse and human results, we constructed a cross-species mapping between homologous cortical and subcortical regions based on published anatomical and functional homologies. We first relied on the homology framework reported in a prior cross-species connectivity study (Balsters et al. [29]), specifically using the set of cortical and limbic regions listed in Supplementary Table 1 of that work, which provides literature-supported correspondences between human cortical areas and their mouse counterparts. This mapping includes medial prefrontal (areas 25, 32pl, and 24), retrosplenial cortex, orbitofrontal cortex (area 13), basolateral amygdala, anterior and posterior hippocampus, primary and secondary somatosensory cortex, primary motor cortex, and temporoparietal association cortex. We did not focus on the original paper’s striatal analyses and instead used the homology table solely as a reference for cross-species cortical and limbic correspondence.

Because thalamic contributions were prominent in one of the LC-derived spatial modes, we extended this mapping to include higher-order and sensory thalamic nuclei. In particular, we incorporated established homologies between the primate pulvinar and the rodent lateral posterior (LP) nucleus, as well as between ventral posterior medial (VPM), ventral posterior lateral (VPL), and ventral lateral anterior (VAL) nuclei across species, based on anatomical and functional evidence from prior comparative and tracing studies [53], [54], [55], [56], [57]. Using this information, we assembled a curated mapping table linking human ROIs from the Harvard–Oxford cortical and subcortical atlases (2 mm, maximum probability threshold of 50%) and the Morel thalamic atlas (2 mm) to corresponding mouse regions from the Allen Brain Atlas. The complete mouse-to-human homology table used in this study is provided in Supplementary Table 1.

Mouse LC-derived SVD maps were summarized at the ROI level by averaging values across anatomically matched mouse regions. These ROI-level values were then transferred to human space by assigning the averaged mouse-derived map weights to the corresponding human ROIs. When multiple mouse regions mapped onto a single human ROI, values were averaged. This procedure yielded homologous human representations of the mouse LC spatial modes, both as ROI-wise summaries and as reconstructed NIfTI volumes in standard MNI space, which were subsequently used for all cross-species analyses.

##### 6.3.5 fMRI – Sleep PSG Dataset Processing and Spatial Regression

For the human sleep–dream dataset, we re-used the publicly available fMRI and polysomnography recordings originally acquired for dream-content decoding during early NREM sleep (Horikawa et al., [28]). Functional data were collected on a 3T scanner using T2*-weighted gradient-EPI (TR = 3000 ms, TE = 30 ms, flip angle = 80°, FOV = 192 × 192 mm, voxel size = 3 × 3 × 3 mm, 50 slices, no slice gap). During continuous scanning, participants repeatedly fell asleep and were awakened by name-calling, after which they provided a brief verbal report; PSG (EEG/EOG/EMG/ECG) was acquired simultaneously and used for sleep staging. In our analyses, we focused on NREM stage 1–2 segments preceding awakenings, and the wake-to-sleep segments following awakenings, capturing the period of wakefulness induced by the call and the subsequent return to NREM. fMRI runs were processed in FSL using a standard pipeline: motion correction (MCFLIRT), spatial smoothing (Gaussian kernel, FWHM = 5 mm), and temporal detrending with a high-pass filter (cutoff = 100 s). Each run was then decomposed with ICA (30 components), and artifactual components (motion/physiological/scanner-related) were manually identified and removed via linear regression, together with residual motion-related variance. To enable direct comparison with the mouse sleep dataset, we used the same signature-expression (spatial regression) approach. SVD-derived LC spatial maps (Map_LC1_ and Map_LC2_ obtained from optogenetic LC stimulation, transformed to human template space as described above) were registered to each run’s native space using an affine (12-DOF) FLIRT transform computed between the MNI template and the first EPI volume, and the same transform was applied to each map. Within a brain mask, each voxel’s fMRI time series was z-scored across time. For each map and each fMRI volume, we then computed a single regression/similarity value as the normalized dot product between the map’s voxel weights and the z-scored fMRI volume, yielding a continuous time course that tracks moment-to-moment expression of the LC-linked spatial pattern. Sleep-stage annotations were taken from the provided hypnograms. We extracted similarity epochs around NREM→Wake and Wake→NREM transitions (±60 TR; TR = 3 s) and quantified post-transition modulation using the same event-wise AUC framework as in mice: baseline correction, within-epoch z-normalization, integration of |z| over 0–30 s, and comparison to an empirical null built from randomly sampled windows from the continuous similarity trace (Kolmogorov–Smirnov test, and Mann–Whitney tests, with Cliff’s δ as an effect size).

##### 6.3.6 Awake Human resting state fMRI – Dataset processing and spatial regression

For the awake human dataset, we analyzed a 7T resting-state fMRI dataset originally reported by Hearne et al. [30] and subsequently analyzed using the neuromodulatory extraction and event-detection framework described by Munn, Müller et al. [25] (N = 59 healthy adults; TR = 586 ms; 2 mm isotropic resolution). In the latter study, the authors extracted time series from key subcortical hubs of the ascending arousal system, including the locus coeruleus (LC) and the basal nucleus of Meynert (BNM), and identified phasic arousal events as rapid, transient increases in the temporal derivative of each signal. We used the LC/BNM time series and event indices provided by Munn, Müller et al., focusing specifically on LC-dominant bursts, defined as events in which LC activity showed a prominent phasic increase while BNM activity showed no concomitant increase or was reduced. The dataset consisted of fMRI data parcellated by the original authors into cortical and subcortical regions using a 2 mm Harvard–Oxford atlas, complemented by a Morel atlas for thalamic subdivisions. All analyses were therefore performed directly in ROI space, without reverting to voxel-level data. To enable direct comparison with the other datasets in this study, we projected the SVD-derived LC spatial maps obtained from optogenetic LC stimulation (Map_LC1_ and Map_LC2_) into the same ROI space. This was achieved by matching template ROIs to the corresponding Harvard–Oxford cortical regions and Morel thalamic nuclei, yielding LC-weighted ROI templates compatible with the parcellated human fMRI time series. Map-expression time courses were computed using the same spatial regression framework applied throughout the study. Specifically, ROI time series were z-scored across time, and for each map a single similarity value per time point was obtained as the normalized dot product between the template weights and the z-scored ROI activity vector. This resulted in continuous time series reflecting the moment-to-moment expression of LC-related spatial patterns. We then used the LC phasic arousal events indices provided by the authors to extract event-centered windows from the map-expression time courses. For each event type, symmetric time windows around each peak were extracted, z-scored within event to control for amplitude differences across subjects and events and averaged to obtain mean and SEM time courses per map-expression. Event-wise modulation of map expression was quantified using a baseline-corrected area-under-the-curve (AUC) metric. Real AUC distributions were compared against a subject-matched empirical null distribution obtained from randomly sampled windows of identical duration using Kolmogorov–Smirnov and Mann–Whitney U tests, with Cliff’s δ reported as an effect size.

## Functional connectivity analysis

To assess how neuromodulatory activity influences large-scale functional coupling, we quantified subject-level variations in functional connectivity and related them to LC-derived brain-state expression. Because the first two SVD modes jointly describe complementary components of the same LC-related pattern, we summarized map-expression using the root-mean-square of their combined magnitude 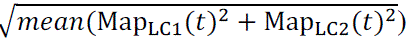, which captures the overall strength of LC-related activity irrespective of the relative contribution of each mode. Global functional connectivity was computed for each subject using the full set of cortical ROIs. ROI time series were z-scored across time, pairwise Pearson correlations were computed, and connectivity strength was summarized as the mean Fisher z-transformed correlation across the upper triangle of the correlation matrix. Finally, to test whether neuromodulatory state expression was associated with large-scale network organization, subject-level LC-related map-expression strength was related to global functional connectivity using Pearson correlation. Confidence intervals for the regression were estimated using non-parametric bootstrap resampling.

## Supporting information

Supplementary Doc 1

## Acknowledgments

We acknowledge the resources and expertise provided by the CIBM Center for biomedical Imaging. We are grateful to the veterinary staff, Jocelin Grosse and Estelle Gerossier, for their support during the urethane fMRI scanning sessions. Language editing assistance was provided by a large language model-based tool; the authors retain full responsibility for the content and its interpretation.

## Author contributions

Conceptualization: VZ, AL

Methodology: FB, GF, DW, LMJF, BRM

Investigation: FB, GF

Visualization: FB, GF

Supervision: VZ, AL, JMS

Writing—original draft: FB, VZ

Writing—review & editing: FB, GF, BRM, DW, JMS, LMJF, AL, VZ

## Data and materials availability

The data supporting the findings of this study will be made publicly available upon publication. Prior to publication, access may be granted upon reasonable request by contacting the corresponding authors. Detailed references to the original sources of all datasets are provided in Supplementary Doc1. All custom analysis code is available on GitHub at: https://github.com/francescabarce/lc-fmri-state-transitions

## Supplementary Materials

**1.** Supplementary Doc 1

