## Supplementary Doc 1 for "A Conserved Locus Coeruleus fMRI Signature of Brain-State Transitions across Sleep, Anesthesia, and Wakefulness"

Francesca Barcellini *et al.*

**This PDF file includes:**

Fig. S1
Table S1
Table S2

**Fig. S2**

**BNM-dominant events do not engage the LC-derived network signature**

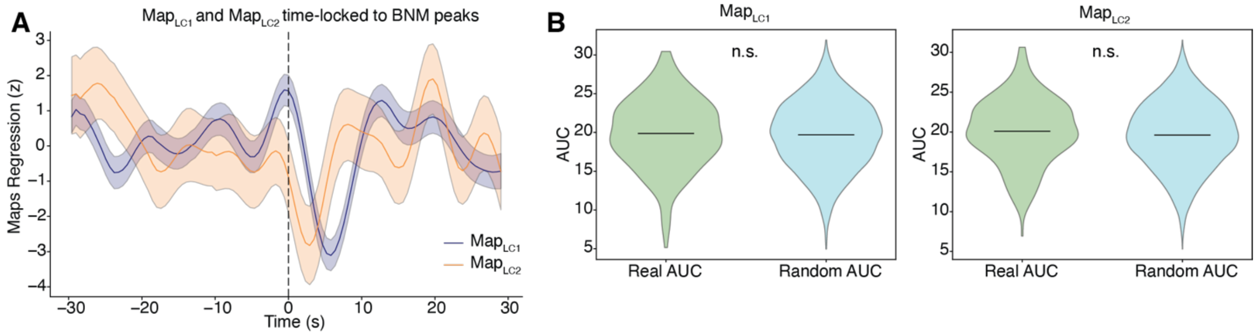


To assess the specificity of the LC-derived network signature for noradrenergic activation, we analyzed fMRI responses time-locked to basal nucleus of Meynert (BNM) peaks occurring in the absence of concurrent LC activation.

**(A)** Event-aligned regression time courses for Map$_{LC1}$ and Map$_{LC2}$ show no consistent, structured modulation around BNM peaks (time 0), indicating a lack of stereotyped LC-like network engagement. Shaded areas denote ± SEM.

**(B)** Event-wise, baseline-corrected AUC distributions for both LC-derived maps did not differ between real BNM events and randomly sampled control windows (n.s., Kolmogorov–Smirnov and Mann–Whitney U tests), confirming that BNM-dominant events do not significantly recruit the LC-associated spatiotemporal signature.

**Table S1**

**Dataset References and Links**

| Dataset type | Data source | Reference |
|---|---|---|
| **Mouse opto - fMRI** | https://zenodo.org/records/7064020#.Yyw2ri8Rpjl | Grimm, C., Duss, S.N., Privitera, M. *et al.* Tonic and burst-like locus coeruleus stimulation distinctly shift network activity across the cortical hierarchy. *Nat Neurosci* **27**, 2167–2177 (2024). https://doi.org/10.1038/s41593-024-01755-8 |
| **Mouse Sleep fMRI** | http://brainliner.jp/data/brainliner/Human_Dream_Decoding | Horikawa, T., Tamaki, M., Miyawaki, Y., & Kamitani, Y. (2013) "Neural decoding of visual imagery during sleep" Science 340, 639-642. http://science.sciencemag.org/content/340/6132/639.long |
| **Human sleep fMRI** | https://www.braindatacenter.cn/datacenter/web/#/dataSet/details?id=1796145515340488706 | Yu Y, Qiu Y, Li G, Zhang K, Bo B, Pei M, Ye J, Thompson GJ, Cang J, Fang F, Feng Y, Duan X, Tong C, Liang Z. Sleep fMRI with simultaneous electrophysiology at 9.4 T in male mice. Nat Commun. 2023 Mar 24;14(1):1651. doi: 10.1038/s41467-023-37352-9. |
| **Human resting state** | https://doi.org/10.5281/zenodo.5315132 | Munn, B.R., Müller, E.J., Wainstein, G. *et al.* The ascending arousal system shapes neural dynamics to mediate awareness of cognitive states. *Nat Commun* **12**, 6016 (2021). https://doi.org/10.1038/s41467-021-26268-x |

### Table S2

### Mouse to Human Homologues region of interest

| Atlas | Index | ROI_Name_Atlas | Mapped_ROI | Filename | Animal_ROI_id | Animal_ROI |
|---|---|---|---|---|---|---|
| HOC | 24 | Frontal Medial Cortex | Area 25 | HOCthr50 | 3 | ILA |
| HOC | 7 | Frontal Pole | Area 32pl | HOCthr50 | 32 | PL |
| HOC | 28 | Cingulate Gyrus, anterior division,Cingulate Gyrus, posterior division | Area 24 | HOCthr50 | 2,4 | ACAd, ACAv |
| HOC | 30 | Precuneous Cortex | Retrosplenial cortex | HOCthr50 | 27,28,29 | RSPd, RSPv,RSPagl |
| HOC | 32 | Frontal Orbital Cortex | Area 13 | HOCthr50 | 24,25 | ORBl |
| HOS | 9,19 | Left Amygdala, Right Amygdala | Basolateral Amygdala | HOSthr50 | 62 | BLA |
| HOS | 8,18 | Left Hippocampus, Right Hippocampus | Anterior Hippocampus | HOSthr50 | 50,51,52,57,55 | CA1,CA2,CA3, ENTl,SUBv, DG |
| HOC | 6 | Precentral Gyrus | M1 | HOCthr50 | 33 | MOp |
| HOC | 16 | Postcentral Gyrus | S1 | HOCthr50 | 9,10,11,12,13,14 | SSp-bfd, SSp-ll, SSp-m, SSp-n, SSp-tr, SSp-ul |
| HOC | 18 | Supramarginal Gyrus, anterior division | S2 (OP1) | HOCthr50 | 15 | SSs |
| HOC | 19 | Supramarginal Gyrus, posterior division | TPJp | HOCthr50 | 1,22 | PTLp |
| MA | 60,152,166 | Ventral Lateral Anterior, Ventral Lateral Posterior Dorsal, Ventral Lateral Posterior Ventral | VLpd,VLpv,VPLp | MA2 | 137 | VAL |
| MA | 23,24 | Ventral Posteriolateral Posterior, Ventral Posteriolateral Anterior | VPLa,VPLp | MA2 | 139,140 | VPL |
| MA | 25 | Ventral Posterior Medial | VPM | MA2 | 139,140 | VPM |

**HOC**: HarvardOxford-Cortical
**HOS**: HarvardOxford-Subcortical
**MA**: MorelAtlasMNI152

**HOCthr50**: HarvardOxford-cort-maxprob-thr50-2mm.nii.gz
**HOSthr50**: HarvardOxford-sub-maxprob-thr50-2mm.nii.gz
**MA2**: morel_2mm.nii.gz
